# Transcriptome-wide splicing quantification in single cells

**DOI:** 10.1101/098517

**Authors:** Yuanhua Huang, Guido Sanguinetti

## Abstract

Single cell RNA-seq (scRNA-seq) has revolutionised our understanding of transcriptome variability, with profound implications both fundamental and translational. While scRNA-seq provides a comprehensive measurement of stochasticity in transcription, the limitations of the technology have prevented its application to dissect variability in RNA processing events such as splicing. Here we present BRIE (Bayesian Regression for Isoform Estimation), a Bayesian hierarchical model which resolves these problems by learning an informative prior distribution from sequence features. We show that BRIE yields reproducible estimates of exon inclusion ratios in single cells and provides an effective tool for differential isoform quantification between scRNA-seq data sets. BRIE therefore expands the scope of scRNA-seq experiments to probe the stochasticity of RNA-processing.

## Background

Next generation sequencing (NGS) technologies have revolutionised our understanding of RNA biology, illustrating both the diversity of the transcriptome and the richness and complexity of the regulatory processes controlling transcription and RNA processing. Recently, efficient RNA amplification techniques have been coupled with NGS to yield transcriptome sequencing protocols to measure the abundance of transcripts within single cells, known as single-cell RNA-seq (scRNA-seq) [1]. scRNA-seq has provided unprecedented opportunities to investigate the stochasticity of transcription and its importance in cellular diversity. Groundbreaking applications of scRNA-seq include the ability to discover novel cell types [2], to study transcriptome stochasticity in response to external signals [3], to enhance cancer research by dissecting tumour heterogeneity [4], to mention but a few. However, such advances have been limited to explore variability between single cells at the gene level, and we know very little about the global variability of RNA splicing between individual cells. Bulk RNA-seq splicing quantification algorithms cannot be easily adapted to the single cell case due to the minute amounts of starting material, low cDNA conversion efficiency and uneven transcript coverage resulting in intrinsically low coverage and potentially high technical noise [5]. This considerably limits the usefulness of scRNA-seq to investigate questions about RNA processing and splicing at the single cell level.

Splicing analysis has been revolutionised by the advent of (bulk) RNA-seq techniques. Early studies [6] quantified splicing by considering junction reads that are uniquely assigned to an inclusion/ exclusion isoform, necessitating very high coverage depth to achieve confident predictions. The situation can be considerably improved by using probabilistic methods based on mixture modelling, an idea that is at the core of standard tools such as Cufflinks [7] and MISO [8]. Nevertheless, low coverage represents a challenge even for probabilistic methods. Recent work has shown that improved predictions at lower coverage can be achieved by incorporating informative prior distributions within probabilistic splicing quantification algorithms, leveraging either aspects of the experimental design, such as time series [9], or auxiliary data sets such as measurements of PolII localisation [10]. Such auxiliary data are not normally available for scRNA-seq data. Nevertheless, recent studies have also demonstrated that splicing (in bulk cells) can be accurately predicted from sequence-derived features [11]. This suggests that overall patterns of read distribution may be associated with specific sequence words, so that one may be able to construct informative prior distributions that may be learned directly from data. Here we introduce the Bayesian Regression for Isoform Estimation (BRIE) method, a statistical model that achieves extremely high sensitivity at low coverage by the use of informative priors learned directly from data via a (latent) regression model. The regression model couples the task of splicing quantification across different genes, allowing a statistical transfer of information from well-covered genes to lower covered genes, achieving considerable robustness to noise in low coverage.

BRIE model has been implemented as a standard Python package, which is freely available at 

~~~
http://github.com/huangyh09/brie.
~~~

 All scripts to replicate the results in this paper are also included in the repository.

## Results and discussion

### High level model description

Figure 1 presents a schematic illustration of BRIE (see Methods for precise definitions and details of the estimation procedure). The bottom part of the figure represents the standard mixture model approach to isoform estimation introduced in MISO [8] and Cufflinks [7], where reads are associated to a latent, multinomially distributed isoform identity variable (see Methods for a self-contained review of mixtures of isoforms models). This module takes as input the scRNA-seq data (aligned reads) and forms the likelihood of our Bayesian model. The multinomial identity variables are the assigned an informative prior in the form of a regression model (top half of Figure 1), where the prior probability of inclusion ratios is regressed against sequence-derived features. Crucially, the regression parameters are shared across all genes and can be learned across multiple single cells, thus regularising the task and enabling robust predictions in the face of very low coverage. In the Methods and Supplementary Material we give details of the features used. While the class of regression models we employ is different from the neural networks of [11], they still provide a highly accurate supervised learning predictor of splicing on bulk RNA-seq data sets. Fig S1 shows that the Bayesian regression approach of BRIE can achieve a Pearson R in excess of 0.8 on test sets, validating our choice of model within BRIE.

**Figure 1.**
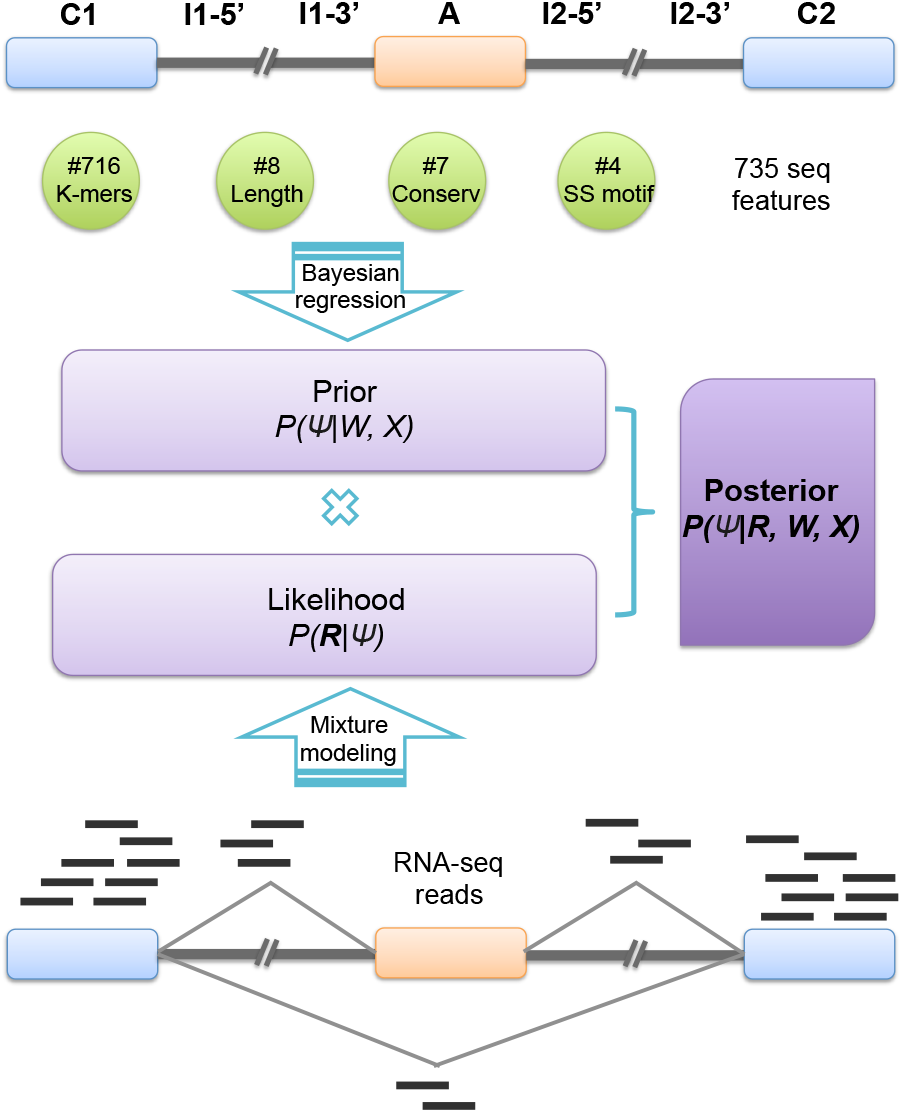
A cartoon of the BRIE method for isoform estimation. BRIE combines a likelihood computed from RNA-seq data (bottom part) and an informative prior distribution learned from 735 sequence-derived features (top).

This architecture effectively enables BRIE to simultaneously trade-off two tasks: in the absence of data (drop-out genes), the informative prior provides a way of imputing missing data, while for highly covered genes the likelihood term dominates, returning a mixture-model quantification. For intermediate levels of coverage, BRIE uses Bayes’s theorem to trade off imputation and quantification.

### Benchmarking BRIE on simulated data

To assess the improvement in isoform quantification afforded by BRIE’s informative prior, we simulated RNA-seq reads for 11,478 human exon-skipping events, and a correlated feature to learn prior (see details in Methods, and Supp. Fig S2). As we are interested in quantifying the effects of an informative prior, we compare BRIE with similar methods developed for bulk RNA-seq: MISO v0.5.3 [8], one of the first and still very widely used probabilistic methods, DICE-seq v0.2.6 [9], a modification of MISO using informative priors (for multiple time points). For completeness, we also compare with Kallisto [12], which was recently proposed as one of the most computationally efficient and robust quantification tools. To simulate the effect of the regression prior, we introduced an auxiliary variable with correlation 0.8 with the desired inclusion ratios (the correlation value was chosen to match the empirical performance of BRIE’s regression prior on bulk RNA-seq data in Supp. Fig S1). We also consider the case when BRIE’s auxiliary variable is uncorrelated with the inclusion ratio (denoted as BRIE.Null) as a control. Thanks to the informative prior, BRIE can also provide an imputation for drop-out transcripts (see below), which other methods cannot; in order to maintain the simulation fair, we did not include results on drop-out genes.

In the simulation, we set different coverage levels, RPK (reads per kilo-base) ranging from 25 to 400. Figure 2 clearly shows that the use of an informative prior can bring very substantial performance improvements at low coverage. At the lowest RPK level, BRIE achieves a gain of almost 20% in correlation between estimates and ground truth. Furthermore, this accuracy level is essentially maintained by BRIE at all coverage values. Interestingly, BRIE.Null can still achieve comparable accuracy to other existing methods at all coverage values; therefore, even in cases where an informative prior could not be effectively learned, BRIE’s results would not be worse than using a state-of-the-art bulk RNA-seq method.

**Figure 2.**
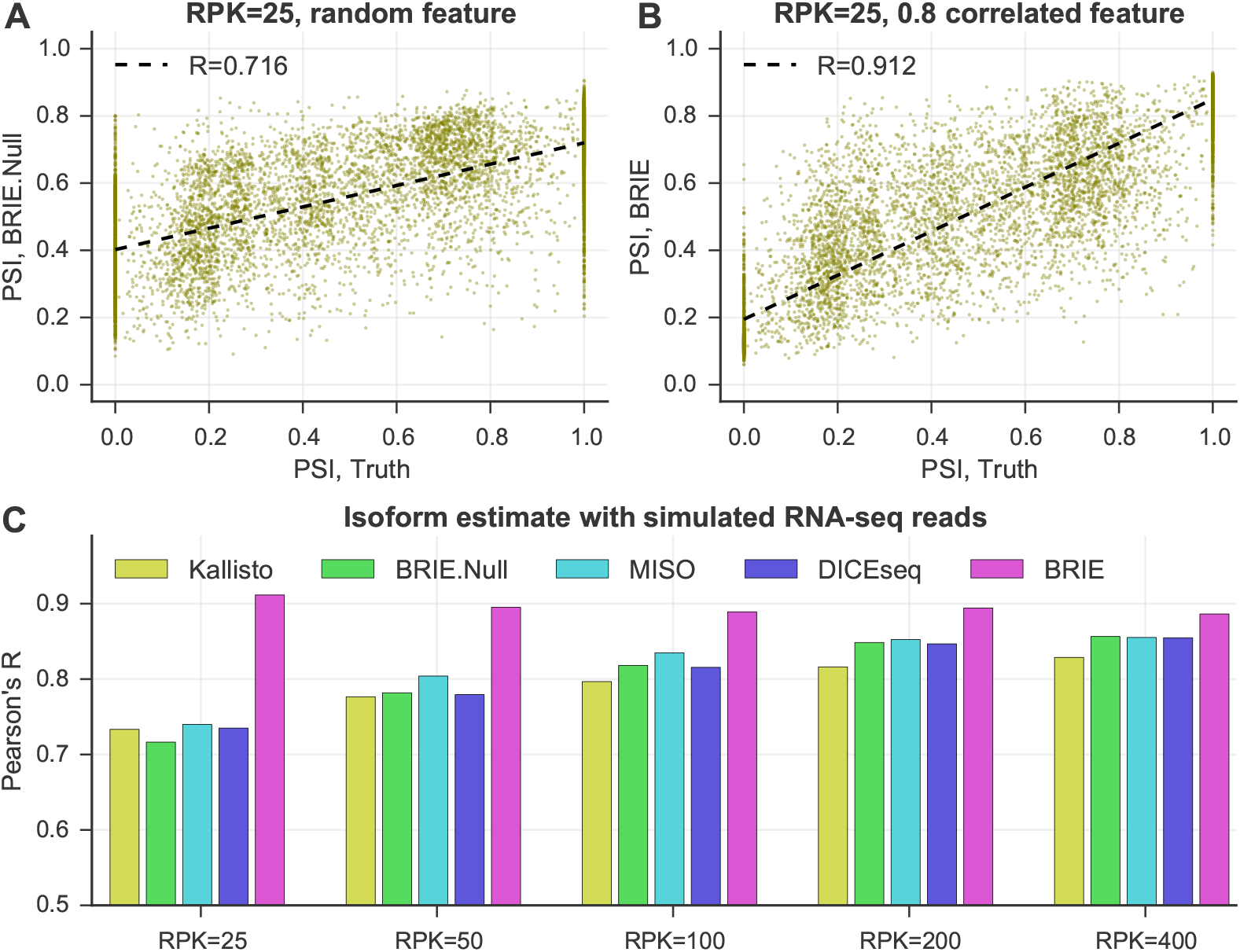
BRIE improves isoform estimates by using an informative prior on simulated data. (A-B) At very low coverage RPK=25, the scatter plot between the estimate of exon inclusion ration by BRIE and the simulation truth. (A) BRIE.Null uses ve random uniform-distributed features to learn prior. (B) BRIE uses one correlated feature with Pearson’s R=0.8 to the truth to learn informative prior. (C) Pearson’s R between truth and estimate by BRIE, BRIE.Null and 3 other methods in di erent coverages.

### Imputation of drop-out in simulation

The informative prior learned by BRIE can also be used to impute isoform usage when there is a dropout, i.e., no reads sequenced for an expressed isoform. In sc-RNAseq experiments, drop-out widely occurs [5], though it is sometimes hard to exactly detect, except for spike-in RNAs. Here, we could coarsely define its upper bound, by counting exon-skipping events expressed in bulk cells but not in a given single cell. In Figure S3, we see that after removing drop-out events, the correlation of expression level between a single cell and bulk cells are dramatically higher on these splicing events.

As BRIE can transfer information from highly expressed gene to lowly expressed genes across multiple cells, we investigated the performance of BRIE in imputing the isoform usage if drop-out happens. Therefore, the expression profile from a bulk RNA-seq library and the drop-out probability profile estimated from 96 HCT116 human cell scRNA-seq libraries ([13], see Fig S4) were used to perform the simulation (see simulation details in Methods). Figure S5 shows that BRIE can produce a good imputation of the isoform usage simply by taking the mean of the informative prior learned from sequence features of the expressed genes (Pearson’s R: 0.6*∼*0.7).

### BRIE yields robust splicing estimates on real data

To assess BRIE’s performance on real scRNA-seq data, we used 96 scRNA-seq libraries from individual HCT116 human cells from the benchmark scRNA-seq study of Wu et al [13] (see Methods for details). Importantly, a bulk RNA-seq data set in the same conditions was also obtained from one million cells. To better explore performance on real data, we expand the set of competing methods to include Cufflinks v2.2.1 [7], RSEM v1.3.0 and the recently proposed single-cell quantification method Census (in Monocle v2.2.0) based on Cufflinks FPKM [14]. Figure 3 shows the results: BRIE clearly outperforms all other methods by a large margin, both in terms of correlation between estimates from different single cells (Fig 3f), and in terms of correlations between estimates from individual single-cells and bulk (Fig 3c). Example scatter plots for both comparisons are given in Fig 3e and 3b, clearly showing very consistent predictions. Notably, the performance of other methods was strongly degraded by the inability to handle the large drop-out rates (see Fig 3a and 3d for DICE-seq, where many estimates of splicing are centred around the uninformative prior value of 0.5). The high correlation between bulk and scRNA-seq predictions is particularly remarkable, as the analysis of the two data sets is not done with a shared prior. Similarly high correlations were found between splicing estimates obtained by BRIE in single cells and estimates from bulk RNA-seq obtained by other methods (Fig S6).

**Figure 3.**
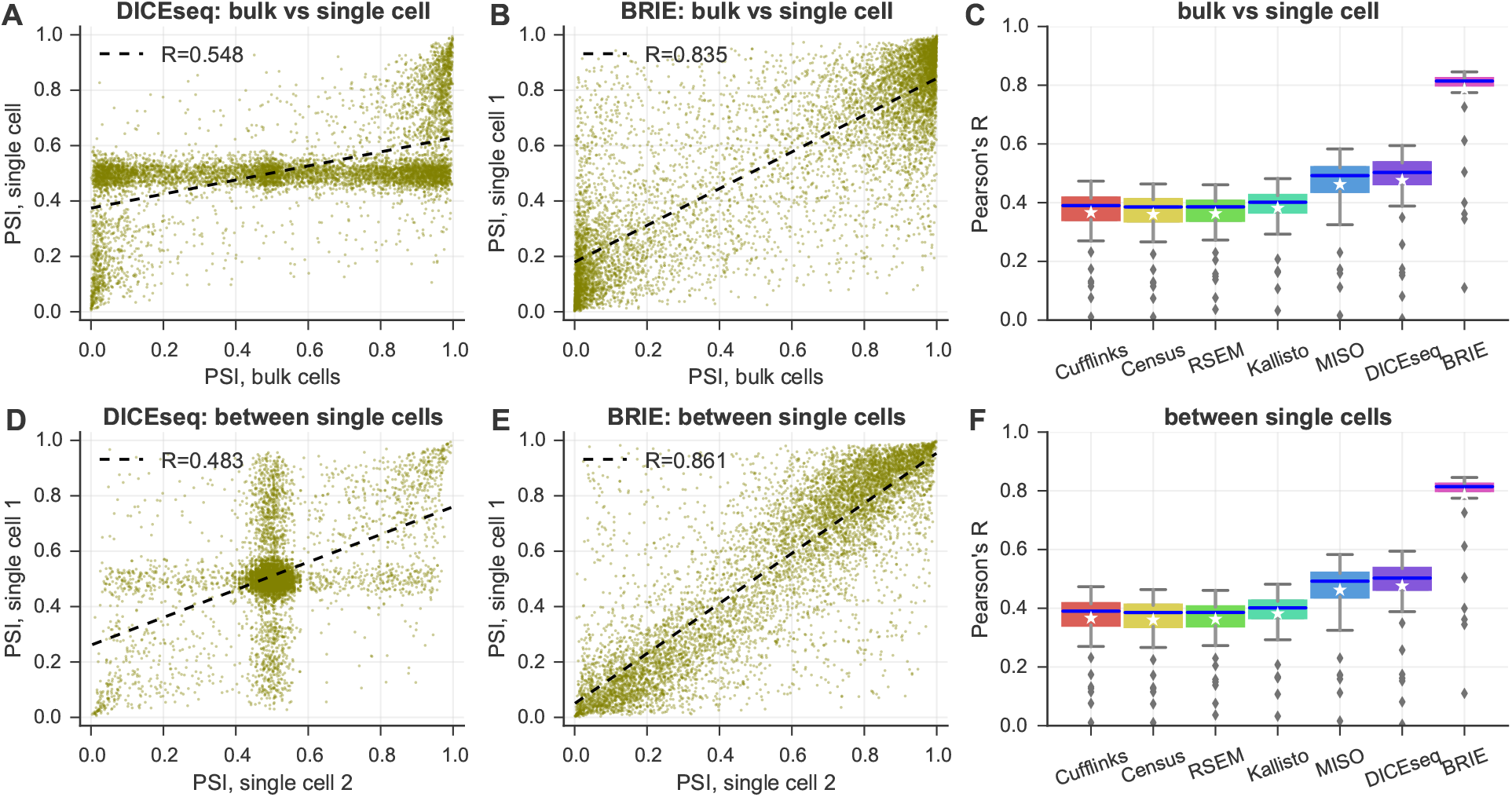
BRIE improves splicing estimates by using sequence features. (A-C) Pearson’s correlation between between bulk and single cells on exon inclusion ratio *Ψ* in HCT116 cells. Scatter plot of *Ψ* estimates by DICEseq (A), or estimated by BRIE (B). Box-plot for all methods (C) in 96 cells. (D-F) Pearson’s correlation between single cell pairs. Scatter plot of estimates by DICEseq (D), or estimated by BRIE (E). Box-plot for all methods (F) in 4,608 cell pairs.

These statistical advantages are reflected in a more effective and confident quantification: considering genes with quantified uncertainty smaller than 0.3 (a threshold adopted e.g. in [15] to select for downstream analysis), Figure S7 shows that BRIE retained 10.9% out of 11,478 genes on average from each single cell (41.1% across all cells), as compared with 3.1% and 5.6% for MISO and DICE-seq, respectively.

### BRIE gives higher sensitivity in differential splicing analyses

BRIE can also be used for differential splicing detection across different data sets. To do so, we compute the evidence ratio (Bayes factor, BF) between a model where the two data sets are treated as replicates (null hypothesis) and an alternative model where the two data sets are treated as separate. We use the Savage-Dickey density-ratio approach and relax it in order to obtain more robust estimates (see Methods). Notice that there are several ways in which differential comparisons could be performed: we could compare groups of cells or individual cells, and we could share the learning of the prior across conditions, or learn separately. All of these options are supported in the BRIE software.

To benchmark the effectiveness of this strategy, we again turned to a simulation study, investigating the ability of BRIE to detect differential splicing as we vary coverage and the extent of the differential effect (see Methods for details of the simulation). This benchmarking is important, as the informative prior might be expected to impede differential quantification. In practice, we see that, for substantial effect sizes (Δ*Ψ* = 0.6), we can detect a substantial fraction of differentially spliced genes already at RPK 50, further improving when the effect size is 0.8 (Fig S8a). We also use the simulation study to explore the effect of different library size on our differential comparisons. We do this by fixing one of the comparison cells to an RPK level. The results shown in Fig S8 b-c demonstrate that BRIE is robust to normalisation issues; this is not surprising, since relative quantification algorithms normally combine normalisation with estimation (see [14] for a discussion of this topic in the scRNA-seq context).

We then moved to investigate the effectiveness of BRIE to detect differential splicing in real cells. To estimate a background level of differential splicing between identical cells, we considered again the 20 single cell HCT116 libraries from Wu et al [13], and compared all possible pairs of cells. Figure 4a shows the fraction of genes called as differentially spliced at different BF thresholds in this control experiment; as we can see, this number is always very small, and around 1% at the normally recommended threshold of BF=10. This level of background calling could be partly attributed to intrinsic stochasticity or to residual physiological variability that was not controlled for in the experiment, such as cell cycle phase. As an additional comparison, we considered two bulk RNA-seq methods for differential splicing, MISO and the recently proposed rMATS [16]. Both methods could only call a negligible number of events, far fewer than the expected number of false positives, confirming that bulk methods are not suitable for scRNA-seq splicing analysis.

**Figure 4.**
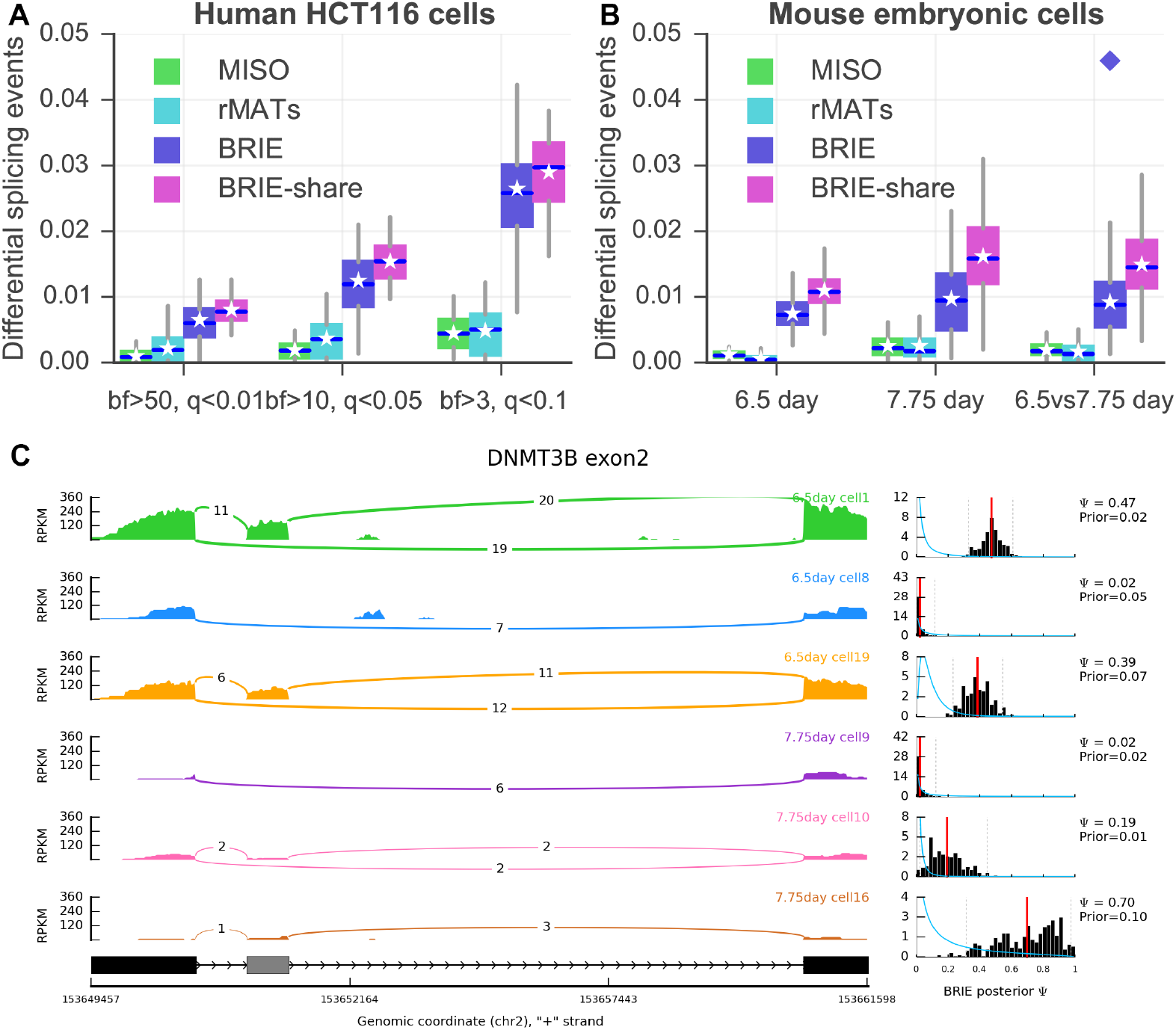
Detection of differential splicing between cells. (A) Percentage of differential splicing events between human HCT116 cells, detected by MISO, rMATS, BRIE and its mode with shared weights (i.e., BRIE.share) with different thresholds. MISO and BRIE use Bayes factor (bf) and rMATS uses false discovery rate (*q* value). (B) Percentage of differential splicing events between mouse early embryonic cells at 6.5 day or 7.75 day. The threshold is *bf* > 10 for MISO and BRIE, and *q* < 0:05 for rMATS. Diamond indicates pooling reads of 20 cells in each group. (C) An example exon-skipping event in DNMT3B in 3 mouse cells at 6.5s days and 3 cells at 7.75days. The left panel is sashimi plot of the reads density and the number of junction reads. The right panel is the prior distribution in blue curve and a histogram of the posterior distribution in black, both learned by BRIE. For the histogram, the red line is the mean and the dash lines are the 95% condence interval.

We then considered a mouse early development scRNA-seq data set [17], and compared the single cell transcriptomic profiles from cells from mouse embryos at 6.5 and 7.75 days. We compared both the profiles of individual cells at the same and different time points; the results are summarised in Figure 4b. Comparing individual cells at 6.5 days yielded approximately 1% of events called as significantly differential (BF≥ 10) at 6.5 days. Comparing this result with our investigation of HCT116 cells suggests that murine cells at 6.5 days are still similar to a homogeneous population, from the splicing point of view. The percentage nearly doubled at 7.75 days, suggesting that differential splicing becomes more widespread at this later stage of differentiation. A similar fraction of exon skipping events were differentially called between cells at 7.75 days and cells at 6.5 days. To define a group of differentiation-associated skipping events, we considered events that we called as differential in at least 10% of 7.75 vs 6.5 comparisons. The resulting 159 events were highly enriched for organelle and intracellular part GO terms (*p <* 0.01) (see Supplementary Table S1 and S2). Figure 4c shows the example of DNMT3B, a regulator of DNA methylation maintenance, which is known to undergo functionally relevant alternative splicing [18]. DNMT3B exhibited differential splicing between 7.75 days and 6.5 days in 153 out of 400 comparisons between individual single cells, clearly highlighting the strong differential inclusion effect. Four more example events, all of which have shown differential splicing in more than 100 pairs of comparisons, are presented in Supplementary Figure S9.

We also directly compared the two groups of cells within a single test (7.75 vs 6.5); this can be easily achieved by assuming a shared splicing ratio *Ψ* across all cells in a condition. Mathematically, this is equivalent to multiplying the likelihood terms associated to each cell, in practice pooling the reads from different cells. While this achieves higher power (see the diamond dot in Fig 4b), it loses the considerable amount of cell-to-cell heterogeneity highlighted by the single-cell analysis. It would be interesting to explore a more refined way of partial pooling within the hierarchical model [19], or to combine BRIE with scRNA-seq clustering approaches which can identify more homogeneous groups of cells [2].

## Conclusions

Our results demonstrate that BRIE can provide a reliable and reproducible method to quantify splicing levels within single cells. Alternative splicing is a major mechanism of regulation of the transcriptome, and splicing analyses within bulk studies have revealed important associations of splicing with disease. Therefore, the ability to quantify alternative splicing in individual cells would considerably expand the relevance of scRNA-seq technology to investigate variations in RNA processing, and its relevance to diseases. We believe the usage of a data-driven informative prior is essential for this task: directly using bulk RNA-seq methods on scRNA-seq is not a viable route due to the limitations of the technology, an observation that was made earlier [1] that our results confirm. Recent work [20] has addressed the issue of *detection* of alternative splicing across a population of single cells, but as far as we are aware BRIE is the first method to be able to *quantify* splicing in individual single cells, and to detect differential splicing between individual ells from scRNA-seq data. We notice that, since BRIE focusses on estimating splicing ratios, it is relatively immune to normalisation issues, since it is essentially a relative quantification method (see [14] for a compelling demonstration of this property of relative quantification methods).

BRIE provides a flexible framework for modelling and, while sequence features are particularly appealing due to their ease of usage and availability, additional side information, such as DNA methylation and chromatin accessibility, could easily be incorporated. Importantly, BRIE is not specific to single-cell RNA-seq technology, and can be of use in any situation where standard quantification is hampered by low coverage.

BRIE’s use of an informative prior enables a smooth trade-off between imputation (at extremely low coverages) and quantification. While this can be a highly effective strategy, it comes at the cost of biasing results at low coverage, potentially introducing some false positives in order to improve the recall of true positives. Another advantage of BRIE’s probabilistic formulation is the ease with which it could be combined with other probabilistic modelling strategies aimed at removing confounders such as cell-cycle stage [21], or at estimating pseudo-time [22].

BRIE cannot be deployed on all scRNA-seq protocols, as it assumes that sequenced reads can be distributed along whole transcripts. Naturally, protocols such as CEL-seq or STRT-seq that bias reads towards the ends of the transcript cannot provide information about exon skipping events that may be very far from the ends of a transcript. We believe that the availability of splicing quantification approaches such as BRIE can therefore be an important consideration in experimental design, particularly at a time when single-cell omic technologies are about to start being more routinely employed.

## Methods

Exon-skipping events annotation

Gene annotations were downloaded from GENCODE human release H22 and mouse release M6. 24,957 and 9,343 exon-skipping events were extracted from protein coding genes on human and mouse, respectively. In order to ensure high quality of the splicing events, we applied 6 constraints following two recent studies [23, 11] for filtering:

1. located on chromosome 1-22 (1-19 for mouse) and X
2. not overlapped by any other AS-exon
3. surrounding introns are no shorter than 100bp
4. length of alternative exon regions between 50 and 450bp
5. with a minimum distance of 500bp from TSS or TTS
6. surrounded by AG-GT, i.e., AG-AS.exon-GT Consequently, 11,478 and 4,549 exon-skipping events from human and mouse respectively were finally used for this study.

### Feature extraction for Bayesian regression

Following Xiong et al [11], we extract predictive sequence features from the following 7 genomic regions for each exon-skipping event (see cartoon in Figure 1a): C1 (constitutive exon 1), I1-5ss (300nt down-stream from the 5’ splice site of intron 1), I1-3ss (300nt upstream from the 3’ splice site of intron1), A (alternative exon), I2-5ss (300nt downstream from the 5’ splice site of intron 2), I2-3ss (300nt upstream from the 3’ splice site of intron 2), C2 (constitutive exon 2).

From these 7 regions, four types of splicing regulatory features are defined. First, 8 length related features are included, i.e., log length of C1, A, C2, I1, I2, and the ratio of the log length of A/I1, A/I2 and I1/I2. Second, the motif strengths of the 4 splice sites, i.e., I1-5’ss, I1-3’ss, I2-5’ss and I2-3’ss, were calculated from mapping each sequence to its averaged position weight matrix. Here, we considered -4nt upstream to +6nt downstream around 5’ss (11nt in total), and from -16nt to 4nt for 3’ss. Third, we also include evolutionary conservation scores for each of the 7 genomic regions, which were calculated by phastCons [24], and are available at the UCSC genome browser. We used the phastCons files in bigWig format with version hg38 for human and mm10 for mouse, where 99 and 59 vertebrate genomes were mapped to the human and mouse genome, respectively. Then the mean conservation scores for the above 7 regions were extracted by using 

~~~
bigWigSummary
~~~

 command-line utility. Lastly, 716 short sequences were extracted from the 7 regions, including 1-2mers for I1-5ss and I2-3ss (20 sequences each), and 1-3mers for C1, I1-3ss, I2-5ss and C2 (84 sequences each), and 1-4mers for A (340 sequences). In total, 735 splicing regulatory features were used to predict the exon inclusion ratio in Bayesian regression.

### RNA-seq data and preprocessing

Bulk RNA-seq libraries for the K562 cell line were produced by the ENCODE project [25], downloaded from Gene Expression Omnibus (GEO: GSE26284); these were used to validate the prediction performance of the splicing regulatory features on bulk RNA-seq (Supplementary Figure S1).

Two single cell RNA-seq data sets were used to validate BRIE model. The first data set is from a benchmark study [13], consisting of 96 single cell RNA-seq libraries from the HCT116 cell line (GEO: GSE51254). These single-cell RNA-seq libraries were prepared with SMART-seq protocol, and have paired-end reads with read length of 125bp. By using a barcode, 48 cells were sequenced per lane, resulting in an average 2.2 million reads per cell. From the same study, two bulk RNA-seq libraries, each with 31.2M reads generated from 1 million HCT116 cells, were also used for comparison. Only reads mapping to alternatively skipped exons and their flanking regions (as described in the previous subsection) were considered.

In order to study differential splicing across different cell types, scRNA-seq data produced by SMART-seq2 protocol from mouse embryo at embryonic day 6.5 and day 7.75 [17] were used. From each of the two groups, 20 individual cells were used, which can be accessed at Array Express (E-MTAB-4079).

All above RNA-seq reads were aligned to the relevant genome reference by HISAT 0.1.6-beta with known splicing junctions.

### Assessing BRIE via a simulation study

There are three simulations conducted to assessing BRIE’s performance in quantifying isoform with low coverages, detecting differential splicing, and imputing splicing in drop-out cases. All synthetic reads were generated by Spanki simulator [26], while we provide Python wraps to easily run the simulations, which is publically available in BRIE GitHub repository.

First, we assessed the robust performance of BRIE in very low coverage on 11,478 human exon-skipping events. We assume that the *Ψ* value follows a 

~~~
logitNormal
~~~

 distribution with mean *μ* = 0 and *σ* = 3, i.e., 

~~~
logit
~~~

(*Ψ*) *∼ N* (0, 3.0), as presented in Figure S2, which is similar to that in ENCODE K562 cell line. Then we set all splicing events at the same sequencing coverage, by fixing its *RPK*, i.e., reads per kilo-base in each experiment. Finally, five different coverage levels are used, including *RPK* = 25 (very low, but comparable to a ly covered gene in a scRNA-seq experiment), *RPK* = 50, *RPK* = 100, *RPK* = 200 and *RPK* = 400.

For the purpose of generating a feature to learn an informative prior, we added Gaussian noise to the output *Ψ* values from the Spanki simulator in its 

~~~
logit
~~~

 format, and ensured a Pearson’s correlation coefficient of 0.8 between the feature and the truth, as shown in Fig S2. This correlation is similar as that achieved by supervised learning in human data set (see Fig S1). By contrast, five uniform-distributed random features are used to learn a Null prior (i.e., random prior), which is named as BRIE.Null.

Second, we tested the power of BRIE in detecting differential splicing events on 400 random mouse exon-skipping events with length ranging from 300bp to 800bp. Eight categories of *Ψ* from 0.1 to 0.9 except 0.5 were equally distributed to the 400 splicing events, and opposite *Ψ* values were assigned to two conditions, e.g., *Ψ*=0.1 in condition 1 and *Ψ*=0.9 in condition 2. Then, the prior is set by the same procedure as the first simulation.

Third, we mimicked the drop-out situation on 11,478 human exon-skipping events, and studied the imputation of BRIE in drop-out cases. We looked at one bulk RNA-seq library and 96 single-cell libraries of HCT116 cell lines [13], and only focus on the splicing events that are expressed in the bulk cells (*FPKM >* 0). We define the drop-out events as those splicing events that are expressed in the bulk cells (*FPKM >* 0) but not in a given single cell (*FPKM* = 0). We further define the drop-out rate of a single cell as the fraction of drop-out events in this cell, and the drop-out probability of a skipping event as the fraction of its drop-out in 96 cells. Both distributions of the drop-out rates and the drop-out probabilities were shown in Fig S4.

Given an expression profile (e.g., FPKM or TPM) *Z* from a bulk library and a profile of drop-out probability calculated from a group of single cells (e.g., the 96 cells here), we simulated the RPK for each isoform (or transcript) as follows. For each isoform *k*, we generate a binary variable *I*_*k*_, i.e., either 0 or 1, following a binomial distribution with mean as its corresponding dropout probability. Then each isoform expression level for the simulated single cell is *αI*_*k*_*Z*_*k*_, where coefficient *α* is included to ensure a given number of total reads. If one wants a different overall drop-out rate, but keep the similarity of the drop-out probability profile, an intercept will be added to the drop-out probability in its 

~~~
logit
~~~

 space. In the simulation of drop-out, the 735 sequence features from real data are used to learn informative prior. We take the mean of the learned prior as the imputed *Ψ* for those drop-out events.

### BRIE model for isoform estimate

Here, we define formally the BRIE statistical model. We consider exon inclusion / exclusion as two different isoforms. We start by reviewing the mixture modelling framework for isoform quantification, introduced in MISO [8]. The likelihood of isoform proportions *Ψ*_*i*_ for observing *N*_*i*_ reads *R*_*i,*1:*N*_*i* in sample (single cell) *i*, can be defined as follows

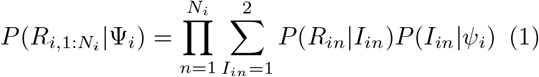

where the latent variable *I*_*in*_ denotes read identity, i.e., the isoform read *n* in cell *i* came from. For bulk RNA-seq methods like MISO [8] or DICEseq [9], the conditional distribution of the read identity *I*_*in*_ *|Ψ*_*i*_ is assumed to be a Multinomial distribution, and the prior distribution over *Ψ*_*i*_ is taken to be an uninformative uniform distribution (suitably adjusted to reflect the potentially different isoform lengths). The precomputed term *P*(*R*_*in*_*|I*_*in*_) encodes the probability of observing a certain read coming from a specific isoform *I*_*in*_. Bulk methods then proceed usually by adopting a Markov-chain Monte Carlo strategy to sample from the posterior distribution of the *Ψ*_*i*_ variables.

BRIE enhances the mixture model approach by combining it with a Bayesian regression module to automatically learn an informative prior distribution by considering sequence features. First, we use a logit transformation of *Ψ*_*i*_, i..e, *y*_*i*_ = 

~~~
logit
~~~

(*Ψ*_*i*_). We then model the transformed exon inclusion ratio *y*_*i*_ as a linear function of a set of *m* covariates *X* ∈ ℝ^*m*^ (here the covariates are the sequence features described previously): *y*_*i*_ = *W*^T^*X* + *ϵ*_*i*_, where *W* is a vector of weights shared by all samples and *ϵ*_*i*_ follows zero-mean Gaussian distribution. All exon skipping events are independently modelled with shared *W* parameters.

Here, we use a conjugate Gaussian prior for the weights, i.e., *W ∼ 𝒩* (0, Λ^−1^), with a common choice of Λ = *λ***I**, for a positive scalar parameter *λ*. Thus, the graphical representation of the full model is shown in Supplementary Figure S10, and the full posterior is as follows (omitting the cell index for simplicity),

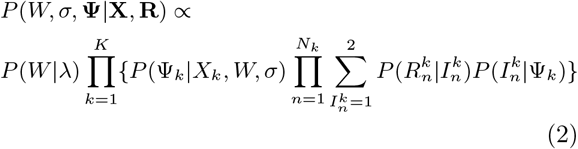

### Inference in BRIE

As shown above, BRIE involves the whole set of exon-skipping events, thus there are thousands of parameters to infer jointly, which can lead to very high computational costs which are not easily distributed. Therefore, we introduce an approximate method to alternately learn *Ψ* and *W*. Also, to alleviate computational burdens, there is an option to merge reads from all cells to learn parameters. For simplicity, we set *λ* empirically, using the value *λ* = 0.1 which gave the best predictive performance on tests on ENCODE data. Then, we collapse *W* and *σ* by taking their expected value in Bayesian regression given a set of *Ψ*, i.e., *W* = (**X**^T^**X** + *σ*^2^Λ)^−1^**X**^T^**Y** and *σ* = 

~~~
std
~~~

(**Y***-W*^T^**X**). At a single exon-skipping event level, we used an adaptive Metropolis-Hastings sampler to sample Ψ, where a univariate Gaussian distribution is used for proposal with adaptive variance, i.e., *η* = 2.38 *\** std(*y*^(1:*m*)^). At this step, we could run short parallel MCMC chains on multiple events to alleviate computational costs, for example *h* = 50 steps if the total iteration is *n*h* = 1000. Pseudocode to sample from the (approximate) posterior distribution of Ψ is given in Algorithm 1. Also, this model supports fixed *W* and *σ*, which can be learned from other data sets, e.g. bulk RNA-seq; then the line 3 and 5 will be turned off in Algorithm 1. The convergence of the sampling is diagnosed by using the Geweke diagnostic *Z* score; in our experiments 1000 burn-in steps appeared to be sufficient in all cases.

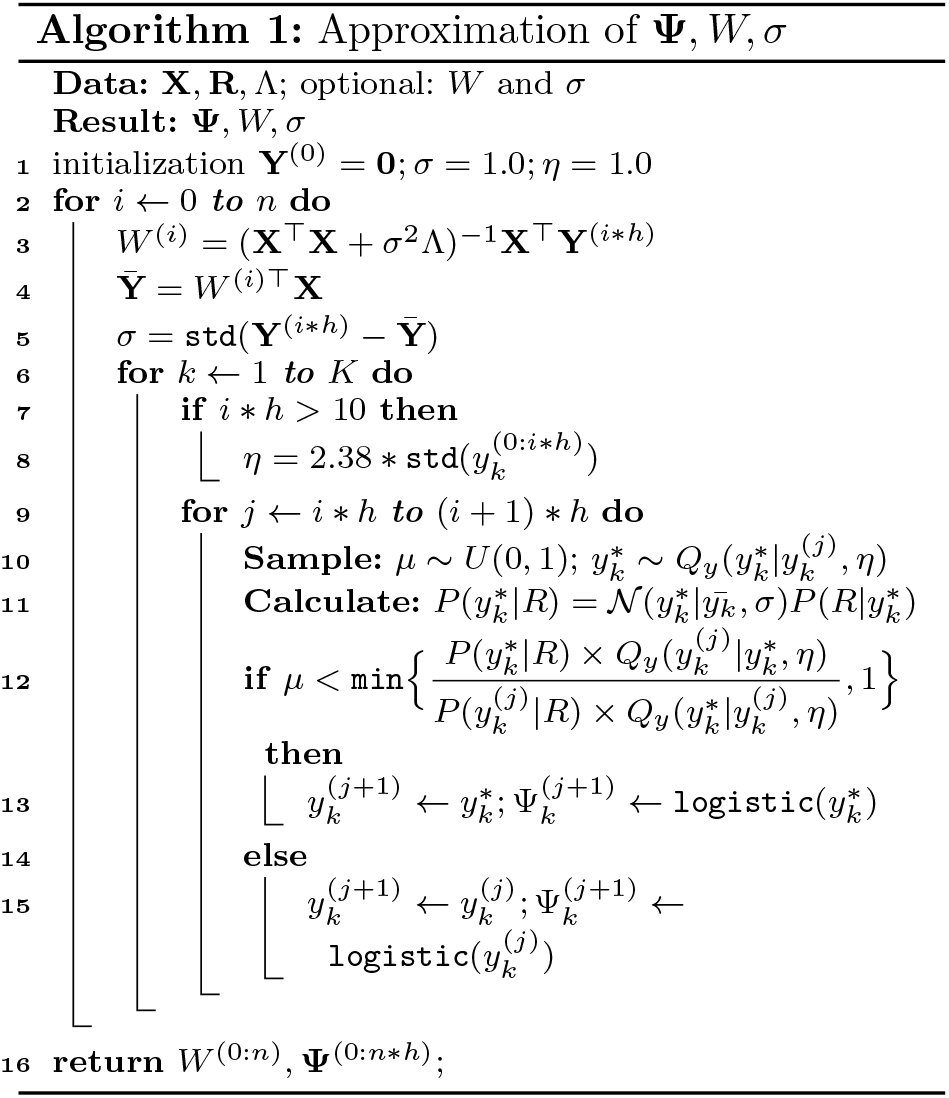

BRIE then outputs an approximate posterior distribution on the *Ψ* values as well as the learned regression weights. BRIE offers functionality to visualise both such posterior distributions as histograms (Fig 4c) and learned weights as heatmaps (Supp. Fig S11 for 19 sequence related features).

### Detection of differential splicing using Bayes factors

The Bayes factor [27] is a posterior odds in favor of a hypothesis relative to another, and is also able to detect whether splicing in two cells or conditions are different or not.

To detect differential splicing between two cells (or cell groups), *A* and *B*, *δ* = Ψ_*A*_ - Ψ_*B*_, we introduce a null hypothesis (*H*_0_) as *δ ≈* 0, and the alternative hypothesis (*H*_1_) as *δ ≉* 0. Here, *D* is the data used to sample the posterior of Ψ in two cells. Then, the Bayes factor in favor of the alternative hypothesis on observing data *D* is defined as follows,

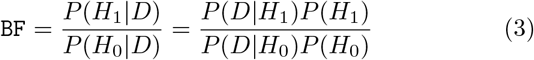

As usual, we assume that both hypotheses have the same prior, i.e., *P*(*H*_1_) = *P*(*H*_0_), and we can clearly see that *P*(*D|H*_0_) = *P*(*D|δ ≈* 0, *H*_1_). Therefore, by taking the Savage-Dickey density ratio [28], we could simplify the calculation of 

~~~
BF
~~~

 as follows,

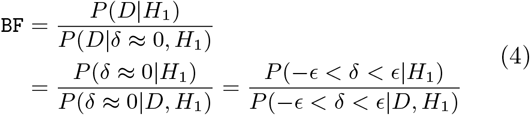

where *ϵ* can be set as 0.05.

As BRIE samples Ψ_*A*_ and Ψ_*B*_ following their posteriors, the distribution of *P*(*δ|D, H*_1_) is readily to approximate by empirically re-sampling Ψ_*A*_ *-* Ψ_*B*_. With a set of re-sampled *δ*_1:*M*_, we take the proportion of *|δ*_*i*_*| < E* as the posterior probability *P* (−*ϵ < δ < ϵ| D, H*_1_). Similarly, we could sample a set of 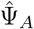 and 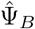 following their prior distributions, and use the same procedure to approximate the prior probability *P*(−*ϵ < δ < ϵ|H*_1_). In the case of comparing two cell groups, one can multiply the individual likelihoods (with shared *Ψ* values); this however is equivalent to pooling reads across different cells, and will loose the quantification of cell-to-cell heterogeneity.

## Abbreviations

scRNA-seq: single-cell RNA-seq; BF: Bayes factor; RPK: reads per kilo-base; MCMC: Markov chain Monte Carlo.

## Author’s contributions

All the authors conceived the study, carried out the data analysis, and wrote the paper. YH developed the software. All authors read and approved the final manuscript.

## Acknowledgements

We thank Chris Ponting and Jean Beggs for useful feedback on an early draft of the manuscript. G.S. acknowledges support from the European Research Council under grant MLCS306999. Y.H. is supported by the University of Edinburgh through a Principal Career Development scholarship.

## Additional Files

Additional file 1 — Figures S1–S11.

Additional file 2 — Table S1-S2.

## References

1. Grün, D., van Oudenaarden, A.: Design and analysis of single-cell sequencing experiments. Cell 163(4), 799–810 (2015)

2. Grün, D., Lyubimova, A., Kester, L., Wiebrands, K., Basak, O., Sasaki, N., Clevers, H., van Oudenaarden, A.: Single-cell messenger RNA sequencing reveals rare intestinal cell types. Nature 525(7568), 251–255 (2015)

3. Shalek, A.K., Satija, R., Shuga, J., Trombetta, J.J., Gennert, D., Lu, D., Chen, P., Gertner, R.S., Gaublomme, J.T., Yosef, N., et al.: Single cell RNA Seq reveals dynamic paracrine control of cellular variation. Nature 510(7505), (2014)

4. Patel, A.P., Tirosh, I., Trombetta, J.J., Shalek, A.K., Gillespie, S.M., Wakimoto, H., Cahill, D.P., Nahed, B.V., Curry, W.T., Martuza, R.L., et al.: Single-cell RNA-seq highlights intratumoral heterogeneity in primary glioblastoma. Science 344(6190), 1396–1401 (2014)

5. Brennecke, P., Anders, S., Kim, J.K., Ko-lodziejczyk, A.A., Zhang, X., Proserpio, V., Baying, B., Benes, V., Teichmann, S.A., Marioni, J.C., et al.: Accounting for technical noise in single-cell RNA-seq experiments. Nature Methods 10(11), 1093–1095 (2013)

6. Wang, E.T., Sandberg, R., Luo, S., Khrebtukova, I., Zhang, L., Mayr, C., Kingsmore, S.F., Schroth, G.P., Burge, C.B.: Alternative isoform regulation in human tissue transcriptomes. Nature 456(7221), 470–476 (2008)

7. Trapnell, C., Williams, B.A., Pertea, G., Mortazavi, A., Kwan, G., Van Baren, M.J., Salzberg, S.L., Wold, B.J., Pachter, L.: Transcript assembly and quantification by RNA-Seq reveals unannotated transcripts and isoform switching during cell differentiation. Nature Biotechnology 28(5), 511–515 (2010)

8. Katz, Y., Wang, E.T., Airoldi, E.M., Burge, C.B.: Analysis and design of RNA sequencing experiments for identifying isoform regulation. Nature Methods 7(12), 1009–1015 (2010)

9. Huang, Y., Sanguinetti, G.: Statistical modeling of isoform splicing dynamics from RNA-seq time series data. Bioinformatics 32(19), 2965–2972 (2016)

10. Liu, P., Sanalkumar, R., Bresnick, E.H., Keleş, S., Dewey, C.N.: Integrative analysis with ChIP-seq advances the limits of transcript quantification from RNA-seq. Genome Research 26(8), 1124–1133 (2016)

11. Xiong, H.Y., Alipanahi, B., Lee, L.J., Bretschneider, H., Merico, D., Yuen, R.K., Hua, Y., Gueroussov, S., Najafabadi, H.S., Hughes, T.R., et al.: The human splicing code reveals new insights into the genetic determinants of disease. Science 347(6218), 1254806 (2015)

12. Bray, N.L., Pimentel, H., Melsted, P., Pachter, L.: Near-optimal probabilistic RNA-seq quantification. Nature Biotechnology 34(5), 525–527 (2016)

13. Wu, A.R., Neff, N.F., Kalisky, T., Dalerba, P., Treutlein, B., Rothenberg, M.E., Mburu, F.M., Mantalas, G.L., Sim, S., Clarke, M.F., et al.: Quantitative assessment of single-cell RNA-sequencing methods. Nature Methods 11(1), 41–46 (2014)

14. Qiu, X., Hill, A., Packer, J., Lin, D., Ma, Y.-A., Trapnell, C.: Single-cell mrna quantification and differential analysis with census. Nature methods 14(3), 309–315 (2017)

15. Barrass, J.D., Reid, J.E., Huang, Y., Hector, R.D., Sanguinetti, G., Beggs, J.D., Granneman, S.: Transcriptome-wide RNA processing kinetics revealed using extremely short 4tU labeling. Genome Biology 16(1), 1 (2015)

16. Shen, S., Park, J.W., Lu, Z.-x., Lin, L., Henry, M.D., Wu, Y.N., Zhou, Q., Xing, Y.: rMATS: robust and flexible detection of differential alternative splicing from replicate RNA-Seq data. Proceedings of the National Academy of Sciences 111(51), 5593–5601 (2014)

17. Scialdone, A., Tanaka, Y., Jawaid, W., Moignard, V., Wilson, N.K., Macaulay, I.C., Marioni, J.C., Göttgens, B.: Resolving early mesoderm diversification through single-cell expression profiling. Nature 535(7611), 284–293 (2016)

18. Duymich, C.E., Charlet, J., Yang, X., Jones, P.A., Liang, G.: DNMT3B isoforms without catalytic activity stimulate gene body methylation as accessory proteins in somatic cells. Nature Communications 7 (2016)

19. Glaus, P., Honkela, A., Rattray, M.: Identifying differentially expressed transcripts from rna-seq data with biological variation. Bioinformatics 28(13), 1721–1728 (2012)

20. Welch, J.D., Hu, Y., Prins, J.F.: Robust detection of alternative splicing in a population of single cells. Nucleic Acids Research 44(8), 73–73 (2016)

21. Buettner, F., Natarajan, K.N., Casale, F.P., Proserpio, V., Scialdone, A., Theis, F.J., Teichmann, S.A., Marioni, J.C., Stegle, O.: Computational analysis of cell-to-cell heterogeneity in single-cell RNA-sequencing data reveals hidden subpopulations of cells. Nature Biotechnology 33(2), 155–160 (2015)

22. Campbell, K., Yau, C.: Ouija: Incorporating prior knowledge in single-cell trajectory learning using bayesian nonlinear factor analysis. bioRxiv, 060442 (2016)

23. Curado, J., Iannone, C., Tilgner, H., Valcárcel, J., Guigó, R.: Promoter-like epigenetic signatures in exons displaying cell type-specific splicing. Genome biology 16(1), 1–16 (2015)

24. Pollard, K.S., Hubisz, M.J., Rosenbloom, K.R., Siepel, A.: Detection of nonneutral substitution rates on mammalian phylogenies. Genome Research 20(1), 110–121 (2010)

25. ENCODE Project Consortium: An integrated encyclopedia of DNA elements in the human genome. Nature 489(7414), 57–74 (2012)

26. Sturgill, D., Malone, J.H., Sun, X., Smith, H.E., Rabinow, L., Samson, M.-L., Oliver, B.: Design of RNA splicing analysis null models for post hoc filtering of Drosophila head RNA-Seq data with the splicing analysis kit (Spanki). BMC Bioinformatics 14(1), 1 (2013)

27. Kass, R.E., Raftery, A.E.: Bayes factors. Journal of the American Statistical Association 90(430), 773–795 (1995)

28. Verdinelli, I., Wasserman, L.: Computing Bayes factors using a generalization of the Savage-Dickey density ratio. Journal of the American Statistical Association 90(430), 614–618 (1995)

